# High-affinity iron and calcium transport pathways are involved in U(VI) uptake in the budding yeast *Saccharomyces cerevisiae*

**DOI:** 10.1101/2021.06.10.447839

**Authors:** Benoît Revel, Patrice Catty, Stéphane Ravanel, Jacques Bourguignon, Claude Alban

**Affiliations:** Univ. Grenoble Alpes, CEA, INRAE, CNRS, IRIG, LPCV, 38000 Grenoble, France; Univ. Grenoble Alpes, CEA, CNRS, IRIG, LCBM, 38000 Grenoble, France

**Keywords:** Yeast, radionuclide, uranium, metal, incorporation, accumulation, mutants

## Abstract

Uranium (U) is a naturally-occurring radionuclide toxic for living organisms that can take it up. To date, the mechanisms of U uptake are far from being understood. Here, we used the yeast *Saccharomyces cerevisiae* as a unicellular eukaryote model to identify U assimilation pathways. Thus, we have identified, for the first time, transport machineries capable of transporting U in a living organism. First, we evidenced a metabolism-dependent U transport in yeast. Then, competition experiments with essential metals allowed us to identify calcium, iron and copper entry pathways as potential routes for U uptake. The analysis of various metal transport mutants revealed that *mid1*Δ, *cch1*Δ and *ftr1*Δ mutants, affected in calcium (Mid1/Cch1 channel) and Fe(III) (Ftr1/Fet3 complex) transport, respectively, exhibited highly reduced U uptake rates and accumulation, demonstrating the implication of these import systems in U uptake. Finally, expression of the *Mid1* gene into the *mid1*Δ mutant restored U uptake levels of the wild type strain, underscoring the central role of the Mid1/Cch1 calcium channel in U absorption process in yeast. Our results also open up the opportunity for rapid screening of U-transporter candidates by functional expression in yeast, before their validation in more complex higher eukaryote model systems.

**Highlights:**

- Living yeast *Saccharomyces cerevisiae* is able to take up U
- Availability of a metabolizable substrate stimulates U uptake
- Calcium, iron and copper inhibit U uptake
- Strains deleted in Mid1/Cch1 calcium channel and Ftr1 iron permease are affected in U uptake
- Expression of *MID1* gene in *mid1*Δ strain restore wild type levels of U uptake

Graphical Abstract

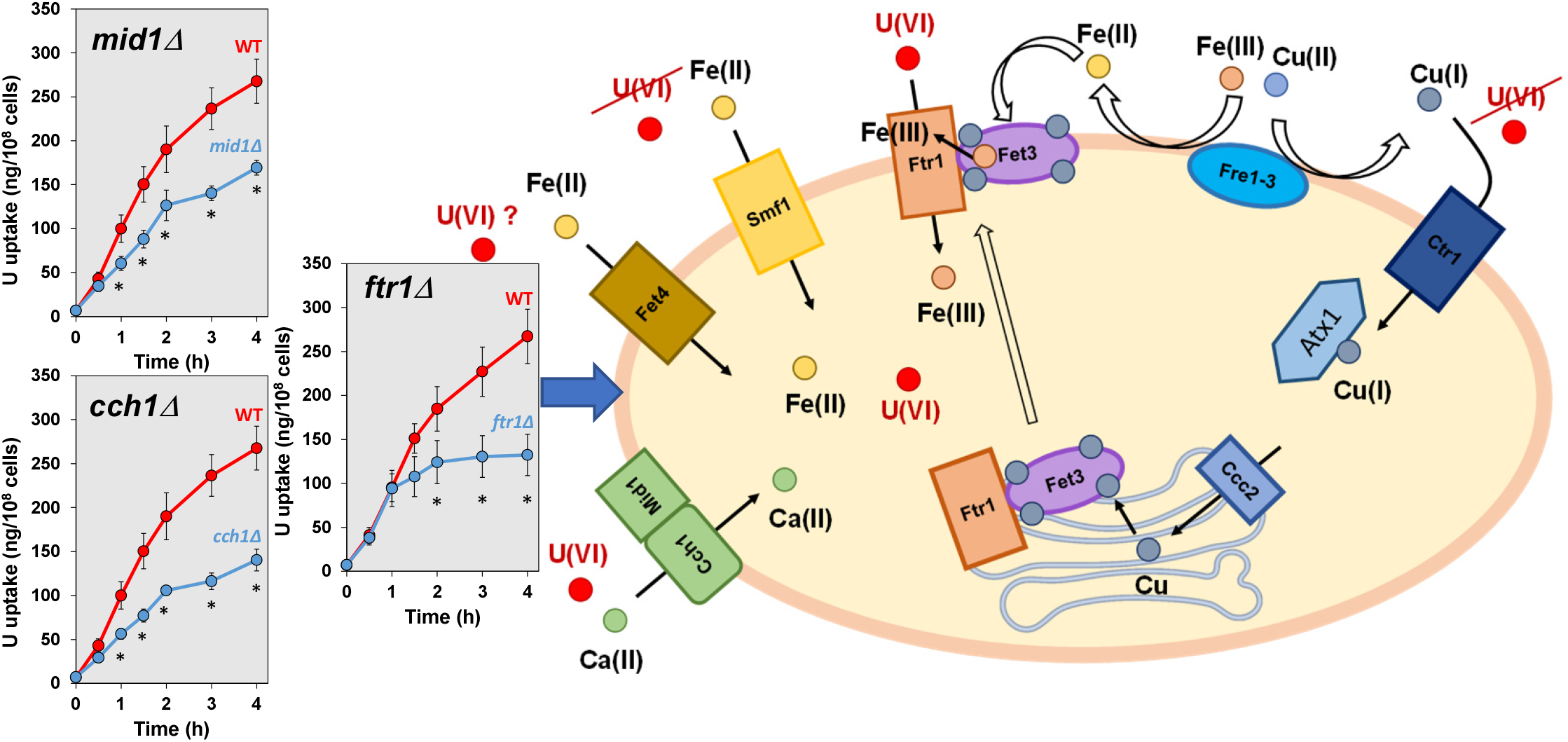

## Introduction

Uranium (U) is a naturally-occurring trace metal element and radionuclide ubiquitous in the Earth crust. It is primarily redistributed in the environment by anthropogenic activities related to U mining and milling industries, civil and nuclear activities, and extensive enrichment of agricultural soils with phosphate fertilizers, which may be significantly contaminated with U. The accumulation of U in soil, water and air can pose potential risks to ecosystems, agrosystems, and ultimately human health through food chain contamination, as the radionuclide has both chemical and radiological effects. Natural U is of low radiotoxicity due to its isotopic composition (>99% ^238^U) but the uranyl cation (UO_2_^2+^) that is prevalent in oxidizing environments is highly chemotoxic (Gao et al., 2019; Ribera et al., 1996). Since U is not an essential element, its uptake and intracellular trafficking depends on existing metal transporters or channels of broad metal selectivity. In particular, cation transporters offer potential transport pathways for toxic metals. These transport systems may differ from a living organism to another. Even if U is known for a long time to interfere with iron, manganese, phosphate and calcium homeostasis in microorganisms, plants and animals, and despite significant efforts to decipher the mechanisms that contribute to the absorption of the radionuclide from the environment, no U entry route in the cell has been identified so far in prokaryotic or eukaryotic cells (Berthet et al., 2018; Gao et al., 2019; Khare et al., 2020) and references therein).

The yeast *Saccharomyces cerevisiae* has long been used as a powerful eukaryotic cell model for studying cell transport systems, in particular those related to trace elements transport and homeostasis (Eide et al., 2005). The usefulness of yeast for genome-wide studies of nutrient homeostasis markedly increased with the completion of the Saccharomyces Genome Deletion Project that resulted in a collection of mutant strains disrupted in most of the approximately 6,000 genes in the yeast genome (Winzeler et al., 1999). This strain collection provides a unique resource for the analysis of gene function in a eukaryotic cell. Moreover, insights from yeast can be easily translated to other organisms, making it an efficient and attractive model system. In many instances, for example, it has been shown that mammalian proteins are capable of functionally replacing yeast proteins, thereby revealing their remarkable functional conservation. Thus, yeast has allowed the identification and molecular understanding of human transporters and has provided insights into diseases that result from defects in ion homeostasis (Cyert and Philpott, 2013; Laliberté and Labbé, 2008; Van Ho et al., 2002). Functional expression of plant genes in yeast also enabled the identification of numerous plant ion transporters before their validation and further characterization *in planta*. This includes, but is not restricted to, the wheat potassium transporter HKT1 (Schachtman and Schroeder, 1994), the Arabidopsis iron transporters IRT1 (Eide et al., 1996), NRAMP1 (Curie et al., 2000), NRAMP3 and NRAMP4 (Thomine et al., 2000), the Arabidopsis MCA1 and MCA2 calcium channels (Nakagawa et al., 2007; Yamanaka et al., 2010) and the Arabidopsis copper transporter COPT1 (Kushnir, 1995). Finally, a systematic approach for transforming yeast into metal hyperaccumulators was recently designed (Sun et al., 2019). This work demonstrated that yeast can be engineered to hyperaccumulate metals by overexpressing and evolving native metal transporters and engineering mechanisms for metal detoxification, allowing to consider the use of this organism as an integral tool for waste treatment processes and recycling.

In this study, we used *S. cerevisiae* to identify pathways for U entry into an eukaryotic cell. U has a strong propensity of biosorption to dead or living yeast cell wall, a process that do not depend on temperature and that has been studied in a bioremediation context (Lu et al., 2013; Shen et al., 2018). Very few studies, however, report on U uptake by yeast cells (Kolhe et al., 2020; Ohnuki et al., 2005; Volesky and May-Phillips, 1995). Here, we developed an efficient and sensitive assay for measuring U uptake in yeast that discriminates U adsorbed on cell surface from internalized U. Uranium uptake was shown to depend on cell metabolism. By using a combination of competition experiments with metal ions and the analysis of metal transport deficient mutants, we showed that calcium, iron and copper assimilation pathways are important routes for U uptake. More precisely, we demonstrated that the high-affinity calcium channel Mid1/Cch1 and the high-affinity iron transporter Ftr1/Fet3 complex are major U importers in *S. cerevisiae*. The potential use of yeast mutants for these transport systems as tools to investigate candidate mammalian and plant U transporters by functional expression, for future human health, phytoremediation and food chain management purposes is discussed.

## Materials and Methods

### Strains and plasmids

Table I lists all yeast strains employed in this study, which were derived from the wild-type *S. cerevisiae* strain BY4742 (MATα, *his3Δ1, leu2Δ0, lys2Δ0, ura3Δ0*) (Brachmann et al., 1998) and were generated by the Genome Deletion Project (Winzeler et al., 1999). All strains were obtained from EUROSCARF.

**Table I:**
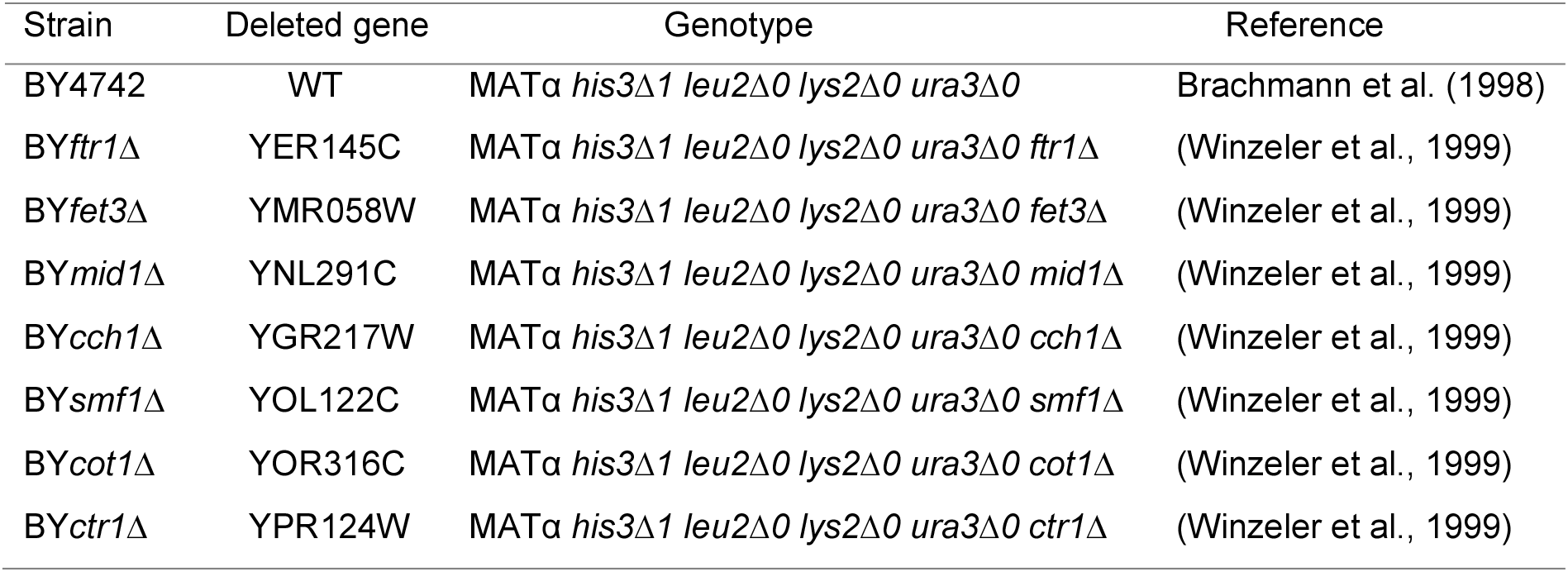
Yeast strains used in this study

Low copy number plasmid YCpMID1 [*URA3 MID1 ARS1 CEN4 amp^r^*] (Iida et al., 2004) was kindly provided by Professor Iida (Tokyo Gakygei University). Empty low copy number plasmid pRS316 [*URA3 ARSH4 CEN6 amp^r^*] (Sikorski and Hieter, 1989) was used as control. High copy number plasmids YEp351-FTR1-myc [*LEU2 FTR1-myc 2 μm-ori amp^r^*], YEp352-FET3-HA [*URA3 FET3-HA 2 μm-ori amp^r^*] and corresponding control empty plasmids YEp351 [*LEU2 2 μm-ori amp^r^*] and YEp352 [*URA3 2 μm-ori amp^r^*] (Stearman et al., 1996) were kindly provided by Doctor Stearman (Indiana University). Plasmids were introduced into yeast strains by using conventional lithium acetate transformation procedure (Kuo and Campbell, 1983). The *Escherichia coli* strain DH5α was used for plasmid amplifications.

### Growth media and culture conditions

Yeast strains were cultured to late-exponential growth phase at 30°C, under continuous stirring at 200 rpm, either in YD medium (10 g/l yeast extract, 20 g/l glucose) or, where indicated, in YG medium (10 g/l yeast extract, 20 ml/l glycerol). Transformed cells were cultured in modified synthetic media SD/Ca100 or SD/Fe0 (6.8 g/l Yeast Nitrogen Base without amino acids; Foremedium Ltd, containing 100 μM CaCl_2_ instead of 900 μM CaCl_2_ or 0 μM FeCl_3_ instead of 1 μM FeCl_3_ in standard mixture, and 20 g/l glucose), supplemented with appropriate amino acids plus adenine (by addition of 0.77 g/l of Complete Supplement Mixture Drop-out formulation minus leucine and/or minus uracil; MPBio). Cells were counted using a LUNA-FL automated cell counter (Logos Biosystems).

### Kinetics of U uptake and accumulation

Cultured yeast cells were harvested by centrifugation at 4000 g for 5 min, washed twice with water and resuspended in 10 mM 2-(N-morpholino)ethanesulfonic acid (MES), pH 5.5 buffer, at a concentration of 2 × 10^8^ cells/ml with or without a carbon source (20 mM glucose, or 3% (v/v) glycerol when cells were grown in YG medium). After a 15 min preincubation at 4°C (in a cold room) or 30°C in a Xtemp climatic chamber (Mechanical Convection Oven XT5116 – IN140) with 30 rpm rotary shaking, uranyl nitrate (UO_2_(NO_3_)_2_.6H_2_O; >98%; Fluka) was added to 10 ml of the cell suspensions to the desired final concentration, from convenient stock solutions. At appropriate intervals, samples (0.5 to 1 ml) were taken and centrifuged at 14 000 g for 2 min. Cell pellets were then washed twice with 1ml 10 mM sodium carbonate. NaN_3_ inhibitor or other metals used in competition experiments were added in the uptake assay at the beginning of the preincubation period. Cell-free controls were conducted using the same procedure to assure that the amount of U precipitating in the reaction medium or adsorbed on the tube wall was negligible under our assay conditions, confirming that U removal resulted exclusively from biotic interaction.

### Determination of cell viability

Yeast viability kit based on fluorescein diacetate/propidium iodide staining method was used to analyze the cell concentration and viability of yeast samples with the automated fluorescence cell counter LUNA-FL, following the manufacturer’s recommendations (Logos Biosystems). In all experiments performed in this study it was found that cell viability was not significantly affected at the end of the absorption kinetics of U (Supplemental Figure 1 A-D).

### U quantification by inductively coupled plasma-mass spectrometry (ICP-MS)

Yeast cells were digested at 90°C for 4 hours in 65% (w/v) HNO_3_ (Suprapur, Merck). Mineralized samples were diluted in 0.5% (v/v) HNO_3_ and analyzed using an iCAP RQ quadrupole mass instrument (Thermo Fisher Scientific GmbH, Germany) equipped with a MicroMist U-Series glass concentric nebulizer, a quartz spray chamber cooled at 3°C, a Qnova quartz torch, a nickel sample cone, a nickel skimmer cone with a high-sensitivity insert, and an ASX-560 autosampler (Teledyne CETAC Technologies, USA). ^238^U was analyzed using the standard mode. Concentrations were determined using standard curves prepared from serial dilutions of UO_2_(NO_3_)_2_ and corrected using an internal standard solution containing ^172^Yb added online. Data integration was done using the Qtegra software (Thermo Fisher Scientific GmbH, Germany).

### Statistical analyses

Statistical parameters including the definitions and values of n and SDs are reported in the figures and corresponding figure legends. Non-parametric statistical analysis was performed on data sets analyzed in Fig. 4, 6 and 7, which typically contain small sample sizes (n<10) and do not meet the assumptions of parametric tests (normal distribution and homogeneity of variance, as determined using the Shapiro-Wilk and Fisher tests, respectively). When reporting significance, multiple non-parametric comparisons were performed with the Dunnett’s many-to-one test using the package nparcomp (Konietschke et al., 2015) and the R computing environment (R Development Core Team, 2011). The normal approximation option was used and the confidence level was set at 99.9 %.

## Results

### *S. cerevisiae* is able to take up U

The budding yeast *S. cerevisiae* is known to adsorb large amounts of U at the cell surface. This massive adsorption of U possibly hides a potential U uptake whose existence and *a fortiori* mechanisms are still largely unknown. To analyze the capacity of yeast cells to transport U intracellularly, we measured kinetics of U incorporation by the strain BY4742 at different U concentrations and two different temperatures. In a first set of experiments, yeast cells (2 × 10^8^ cells/ml) were challenged with 10 μM uranyl nitrate in 10 mM MES buffer pH 5.5 in the presence of 20 mM glucose for up to 3 h, at 4°C or 30°C. Cellular U content was measured by ICP-MS after two washing steps with 10 mM sodium carbonate. As shown in Figure 1, more than 70% of U initially added to the reaction medium was recovered immediately in the unwashed cell pellets (time 0 corresponds to the time necessary to separate cells from surrounding buffer by centrifugation after U exposure, *i.e.* less than 5 min). This fast and temperature independent U binding to yeast cells corresponded to adsorption. Indeed, metal cations adsorption on yeast cell wall is reported to be unchanged across the temperature range 4-30°C (Blackwell et al., 1995; White and Gadd, 1987). At 4°C, the level of biosorbed U remained stable throughout the kinetics. After one washing step with sodium carbonate most of U adsorbed at the cell surface was washed out and recovered in the washing solution after centrifugation (Figure 1A). After the second washing step, almost all adsorbed U was eliminated (U measured in cell pellets did not exceed the background value throughout the kinetics, demonstrating the efficiency of the washing procedure). This demonstrated that almost no U enters yeast cells at 4°C. At 30°C, U associated to the cell pellets was only partially washed away with sodium carbonate. Indeed, a time-dependent increase of U incorporated was observed in washed cells (starting from nearly 0 immediately upon addition of U), which correlated with a decrease of U remaining in the incubation medium and in the washing solutions (Figure 1B). Under these experimental conditions, we measured a rate of 70-100 ng U (0.29-0.42 nmol) transported per hour by 10^8^ cells. Altogether, these experiments showed that yeast cells are able to incorporate intracellularly in a temperature dependent manner a portion of U added exogenously after a rapid and massive adsorption at the cell surface.

**Figure 1:**
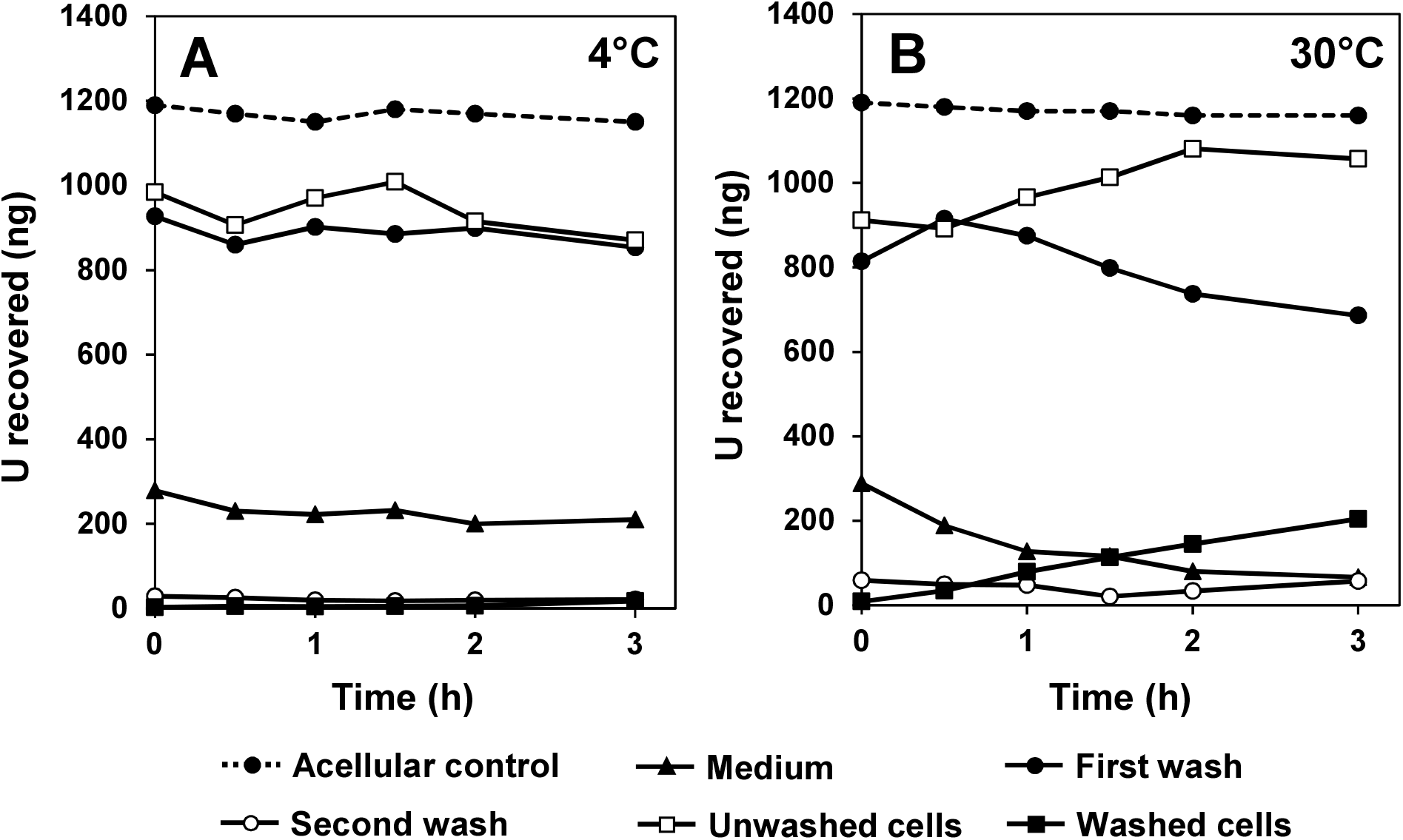
Effect of sodium carbonate washes on unmasking U taken up by U-exposed *S. cerevisiae* BY4742 strain cells. Cells grown at late-log phase in YD medium were suspended at a concentration of 2 × 10^8^ cells/ml in 10 mM MES, pH 5.5 buffer containing 20 mM glucose, and preincubated for 15 min at 4°C (A) or 30°C (B) before addition of 10 μM uranyl nitrate and further incubation at the above-mentioned temperatures. At the time indicated, 0.5 ml aliquots of the cell suspensions were withdrawn and processed as detailed in the Materials and Methods section. U remaining in the incubation medium after centrifugation of the cell suspensions (“Medium”), in the washing solutions after two washes of pelleted cells with 10 mM sodium carbonate (“First wash” and “Second wash”, respectively), and in unwashed and washed cell pellets (“Unwashed cells” and “Washed cells”, respectively), was quantified by ICP-MS. Dotted line indicates U remaining in the incubation medium of an acellular control, after centrifugation (“Acellular control”). A result representative of at least three independent experiments is shown. In all experiments, the U recovery profiles were very similar, although the absolute values varied from an experiment to the other.

### U transport in *S. cerevisiae* requires an active metabolism

To learn more about U transport mechanisms in *S. cerevisiae*, the effect of energy source on U uptake was investigated. To this aim, cultured cells suspended in MES buffer were challenged at 30°C and 4°C with increasing amounts of uranium nitrate from 1 to 10 μM either in the presence or absence of glucose for up to 24h. As already suggested by experiments depicted in Figure 1, a huge difference in U uptake was observed between 30°C and 4°C (Figure 2). At 30°C, an important dose-dependent U influx was observed in the presence of glucose. Under these conditions, U uptake was linear for at least 3h for all U concentrations tested and after 24 h incubation, 30% to 40% of U initially present in the reaction mixture accumulated into the cells (Figure 2). In the absence of glucose, accumulation levels of U were much lower. Interestingly, the ratio between the glucose-dependent and the glucose-independent U accumulation increased with U concentration (ratio around 2 at 1 and 3 μM U (Figure 2A,B); ratio of 6 at 10 μM U (Figure 2C)). U uptake measured at 4°C was very low, but for the three U concentrations tested, a positive effect of the presence of glucose on U uptake was also observed. The transport rate of U was assessed in the presence of glucose at 30°C on 90 min kinetics. As shown in Figure 3, an apparent *V*_max_ of 145±5 ng (0.6 nmol) U incorporated per hour by 10^8^ cells was reached at saturated concentrations above 50 μM and an apparent *K*_m_ for U uptake of 9.0±1.4 μM was calculated from the Michaelis Menten equation (Figure 3). Note that cell viability was checked throughout the experiment and was found to be largely unaffected regardless of the tested condition (Supplemental Figure 1A). The stimulation of U uptake in the presence of glucose might be attributable to metabolism-dependent intracellular U accumulation. The requirement of an active metabolism was also checked using glycerol, an alternate but non-fermentable carbon source, and sodium azide, an inhibitor of the mitochondrial respiratory chain. As shown in Figure 4, similar U transport rates were measured in the presence of glycerol and glucose in the absence of sodium azide. The latter almost completely abolished U uptake, regardless of the energy source used in the assay. Altogether, these results demonstrate that U transport within yeast cells is essentially a metabolism-dependent process that requires a functional respiratory chain able to produce ATP.

**Figure 2:**
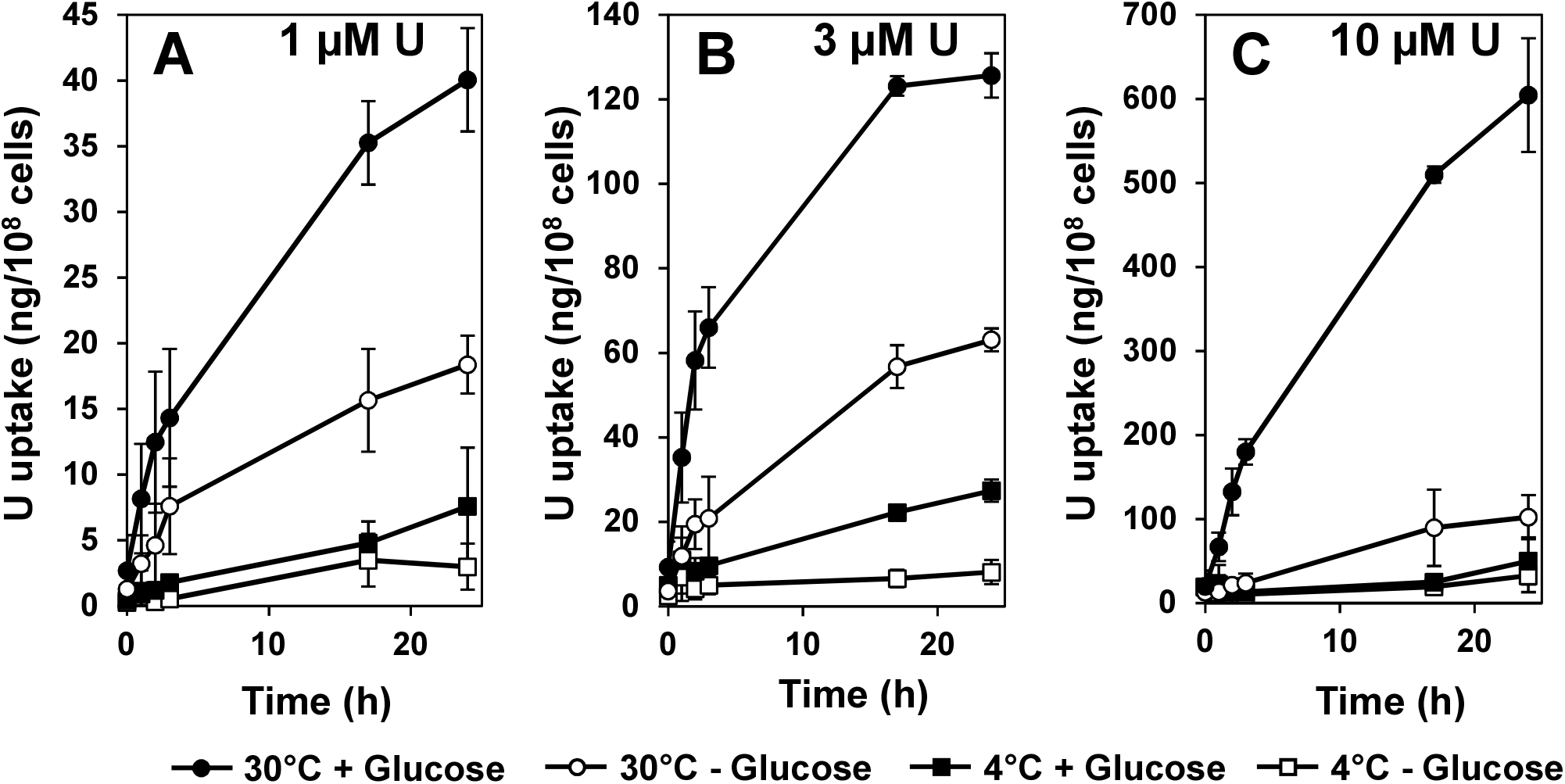
Stimulation of U uptake rate by glucose in *S. cerevisiae* BY4742 strain. Cells grown at late-log phase in YD medium were suspended at a concentration of 2 × 10^8^ cells/ml in 10 mM MES, pH 5.5 buffer containing 20 mM glucose (+ Glucose) or not (- Glucose), and preincubated for 15 min at 4°C or 30°C before addition of 1 μM (A), 3 μM (B) or 10 μM (C) uranyl nitrate and further incubation at the above-mentioned temperatures. At the time indicated, 0.6 ml aliquots of the cell suspensions were taken and processed as detailed in the Materials and Methods section. U present in cell pellets washed twice with 10 mM sodium carbonate was quantified by ICP-MS. Bars = SD; n = 3 independent experiments.

**Figure 3:**
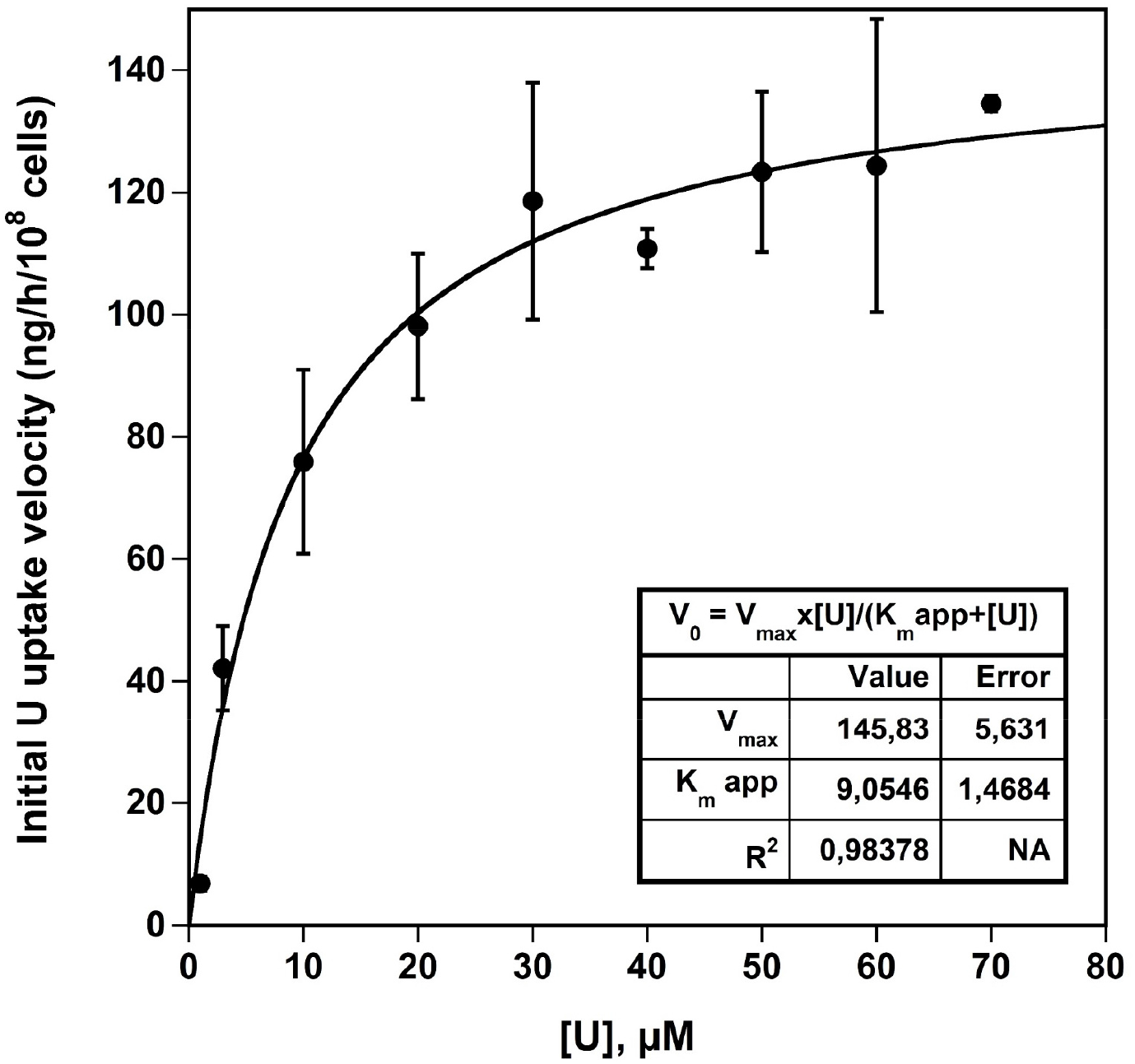
Kinetic parameters determination of U uptake in *S. cerevisiae* BY4742 strain. Cells grown at late-log phase in YD medium were suspended in 10 mM MES, pH 5.5 buffer containing glucose 20 mM, at a concentration of 2 × 10^8^ cells/ml and preincubated for 15 min at 30°C before addition of 1 to 70 μM uranyl nitrate and further incubation at 30°C. At appropriate intervals (from 0 to 90 min), aliquots of the cell suspensions were taken and processed as detailed in the Materials and Methods section. U present in cell pellets washed twice with 10 mM sodium carbonate was quantified by ICP-MS. The initial velocity of U uptake (uptake rate) was then plotted versus U concentration in the reaction medium. Data (dots) were fitted by nonlinear regression to the simple Michaelis-Menten equation (curve) using the Kaleidagraph 4.5 program (Synergy Software, Reading, PA). Bars = SD; n = 2 to 9 independent experiments, for each data point.

**Figure 4:**
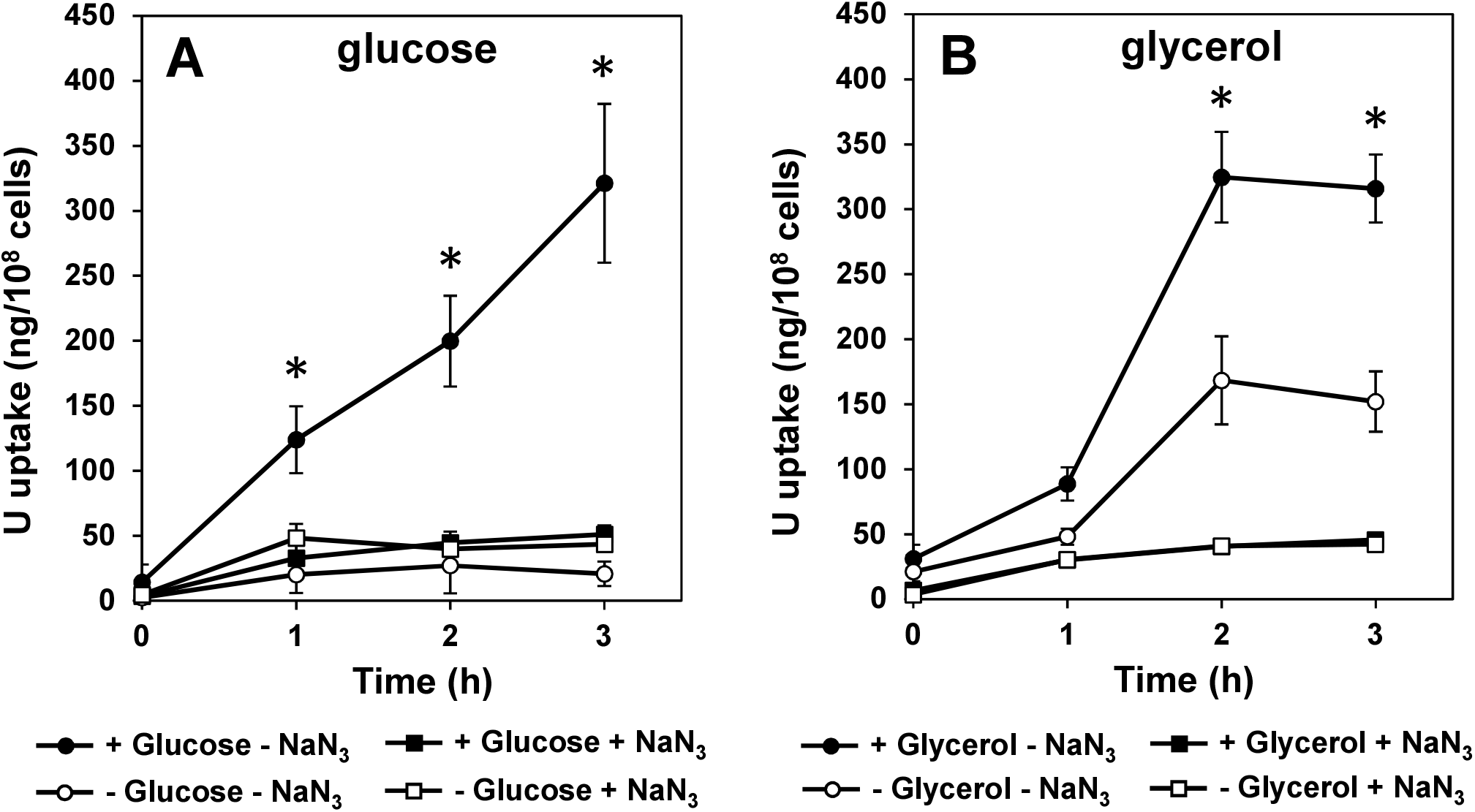
Effect of glycerol as an alternate metabolizable substrate to glucose for U uptake in *S. cerevisiae* BY4742 strain. Cells grown at late-log phase in YD (A) or YG (B) medium were suspended at a concentration of 2 × 10^8^ cells/ml in 10 mM MES, pH 5.5 buffer containing either 20 mM glucose (+ Glucose) or not (−Glucose) (A) or 3% glycerol (+ Glycerol) or not (- Glycerol) (B) and 10 mM NaN_3_(+ NaN_3_) or not (- NaN_3_), and preincubated for 15 min at 30°C before addition of 10 μM uranyl nitrate and further incubation at the same temperature. At the time indicated, 0.6 ml aliquots of the cell suspensions were taken and processed as detailed in the Materials and Methods section. U present in cell pellets washed twice with 10 mM sodium carbonate was quantified by ICP-MS. Bars = SD; n = 3 independent experiments. Statistical significance for the comparison of all other conditions versus + Glucose (or + Glycerol) – NaN_3_ was determined using a non-parametric Dunnett’s test with *P* < 0.001 (*).

### Fe^2+^, Fe^3+^, Cu^2+^ and Ca^2+^ compete with U for transport in *S. cerevisiae*

As a first step toward the identification of U uptake pathways in yeast, its accumulation was measured in the presence of various essential metals whose transport systems are well described. Indeed, since U is not essential for any living organism, its uptake probably depends, at least in part, on these transport systems that, taken together, display a broad metal selectivity, including toxic metals (Cyert and Philpott, 2013; Van Ho et al., 2002). In these competition experiments, U uptake (from 10 μM uranyl nitrate) was challenged with nine essential metals (Fe(III), Cu(II), Fe(II), Ca(II), Mo(VI), Mn(II), Zn(II) Co(II) and Mg(II)) at concentrations varying from 10 μM to 1 mM.

As shown in Figure 5, three metals, namely copper, iron and calcium competed with U uptake. CuSO_4_ (Figure 5A) and FeCl_3_ (Figure 5B) displayed the strongest concentration-dependent inhibitory effect. At 10 μM, both metals had no or limited effect on U uptake. At 50 μM, they caused a 2- and 3-fold reduction of U accumulation after 4h, respectively. The inhibitory effect of these two metal ions was even more marked at 100 μM with a 4-fold reduction of U accumulation after 4 h. If we consider the initial rate of U transport as measured over the first 90 min (see also Figures 2 and 3), it turns out that the low accumulation of U in the presence of Cu(II) and Fe(III) likely came from a reduction in the initial rate of U uptake. The inhibitory effects of Ca(II) (Figure 5C) and Fe(II) (Figure 5D) were different from those of Cu(II) and Fe(III) in our competition assays. They were globally lower and observed at higher concentrations. Indeed, 1 mM of CaCl_2_ or FeSO_4_ were required to induce a nearly 2-fold reduction of U uptake at 4h. Remarkably, Ca(II) and Fe(II) did not seem to significantly affect the rate of U uptake during the first hour of competition (Figure 5 C,D). Finally, Figure 5 E-I shows that, at all the concentrations tested, MgCl_2_, MnCl_2_, ZnSO_4_, Na_2_MoO_4_ and CoCl_2_ had almost no effect on U uptake. Again, cell viability was checked throughout the experiment and was largely unaltered regardless of the condition tested (85-99% viability at the end of the experiment, depending on individual cases) (Supplemental Figure 1C), ruling out the possibility that toxic effects of high concentrations of competing metals were primarily responsible for the reduced U uptake rates and accumulation levels. Taken together, our results identified three potential entry pathways for U in yeast; the calcium, copper and iron uptake routes.

**Figure 5:**
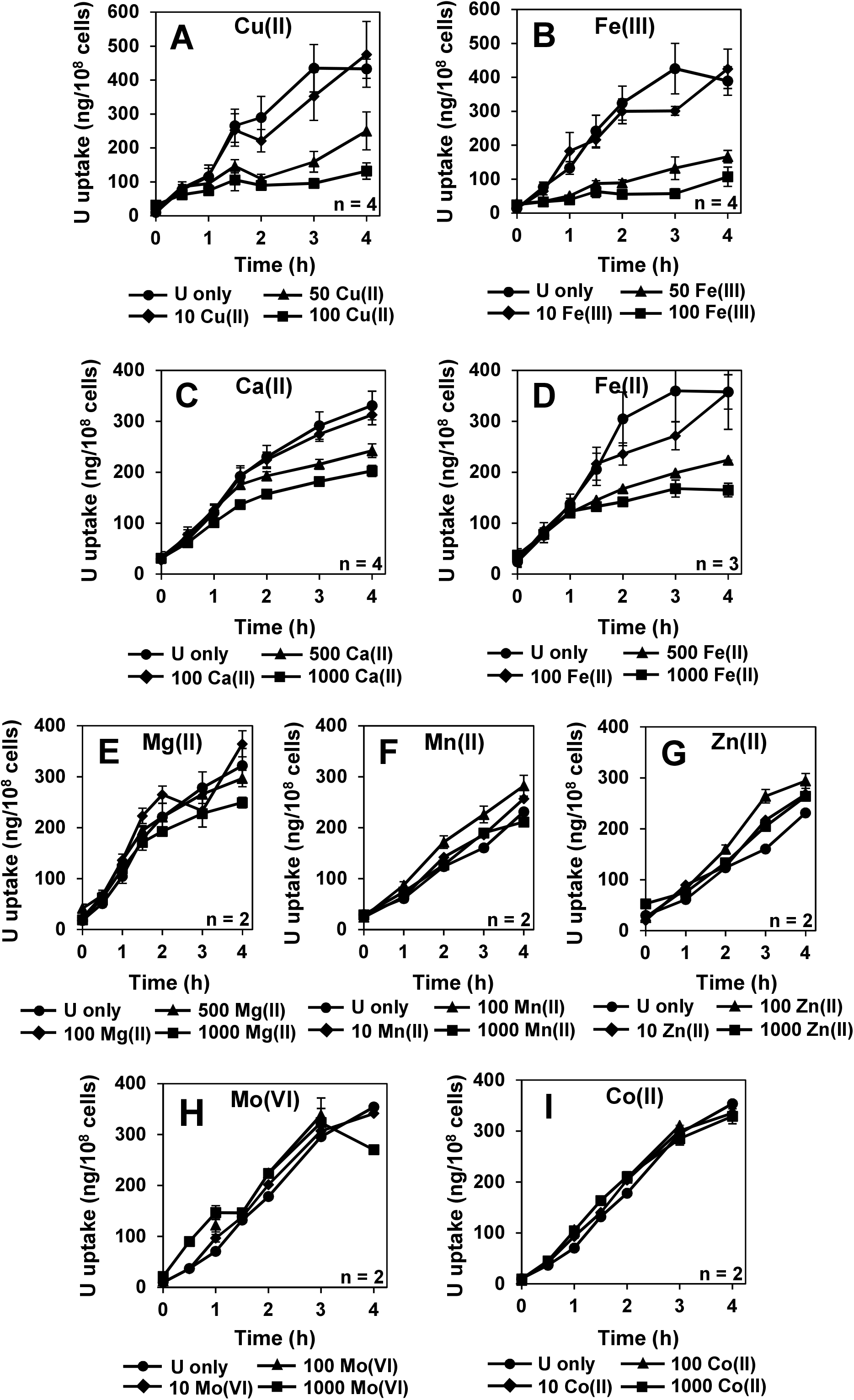
Kinetics of U uptake in *S. cerevisiae* BY4742 strain in the presence of potential essential metal competitors. Cells grown at late-log phase in YD medium were suspended at a concentration of 2 × 10^8^ cells/ml in 10 mM MES, pH 5.5 buffer containing 20 mM glucose and various concentrations (0 to 1000 μM as indicated) of CuSO_4_ (A), FeCl_3_ (B), CaCl_2_ (C), FeSO_4_ (D), MgSO_4_ (E), MnSO_4_ (F), ZnSO_4_ (G), Na_2_MoO_4_ (H) or CoCl_2_ (I). Cell suspensions were then preincubated for 15 min at 30°C before addition of 10 μM uranyl nitrate and further incubation at the same temperature. At the time indicated, 0.5 ml aliquots of the cell suspensions were taken and processed as detailed in the Materials and Methods section. U present in cell pellets washed twice with 10 mM sodium carbonate was quantified by ICP-MS. Bars = SD when these values exceed the dimensions of the symbols; n = 2 to 4 independent experiments, as indicated.

### *S. cerevisiae* iron and calcium transport mutants are affected in U uptake

In order to identify metal transporters involved in U uptake in yeast, a set of mutant strains isogenic of BY4742 and affected in the uptake of various metals were assayed for their capacity to accumulate U. The strains *ftr1*Δ and *fet3*Δ are deleted in the genes coding for the permease Ftr1 and the multi-copper ferrous oxidase Fet3, two components of the high-affinity iron Fe(III) uptake system (Stearman et al., 1996). The strain *smf1*Δ is deleted in the gene coding for Smf1 a high-affinity manganese Mn(II) transporter of the Nramp family that can also transport Fe(II), Cu(II) and other divalent metal ions into the cytosol (Chen et al., 1999; Cohen et al., 2000). The strain *ctr1*Δ is deleted in the gene coding for Ctr1 a high-affinity copper transporter that mediates Cu(I) uptake (Dancis et al., 1994). The strains *cch1*Δ and *mid1*Δ are deleted in the genes coding for the two components of the high-affinity plasma membrane voltage-gated calcium channel Cch1, the pore-forming subunit and Mid1, the positive regulatory subunit (Iida et al., 2004; Iida et al., 2017; Iida et al., 2007; Paidhungat and Garrett, 1997). Finally, the stain *cot1*Δ is deleted in the gene coding for Cot1, a transporter induced in iron-deficient yeast that mediates vacuolar accumulation of zinc and cobalt as well as other metals including toxic ones (Conklin et al., 1992; MacDiarmid et al., 2000).

U uptake kinetics in these seven strains are presented in Figure 6. As shown in Figure 6 A,B,C, U uptake was significantly affected in the *ftr1*Δ, *mid1*Δ and *cch1*Δ strains as compared to the wild type. Note that in the *ftr1*Δ strain, the initial rate of U uptake was not altered as compared to the wild type, the inhibitory effect being only visible after 1 h incubation. Some inhibition of U uptake was detected in the *ctr1*Δ and *fet3*Δ mutants but it was not statistically significant (Figure 6 D,E). In contrast, no difference in U uptake was observed in the *smf1*Δ and *cot1*Δ mutants as compared to their wild type counterpart (Figure 6 F,G). Taken together, these data strongly suggest that yeast cells incorporate U at least through the high-affinity iron transporter Ftr1 and the calcium channel Cch1/Mid1.

**Figure 6:**
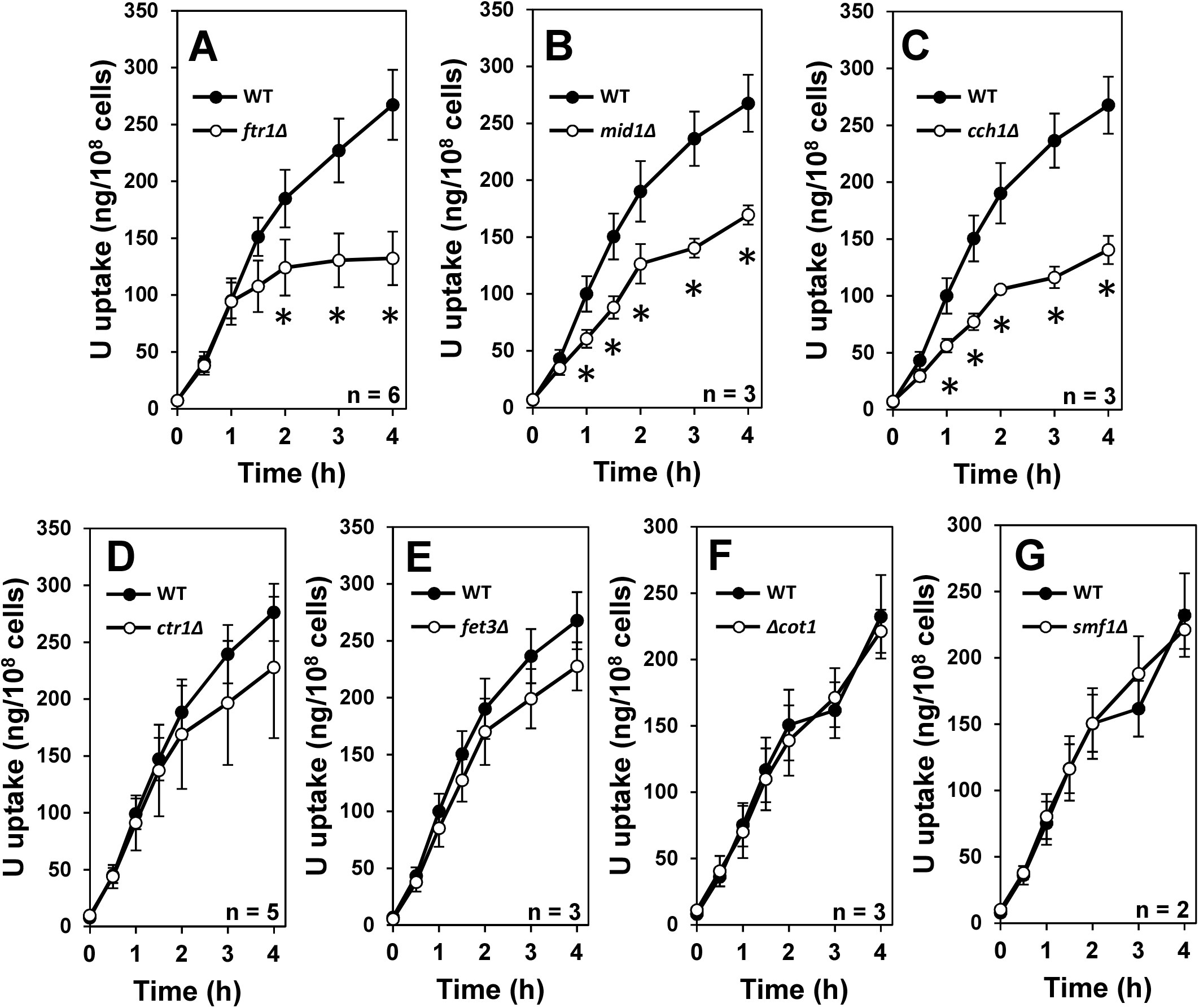
Kinetics of U uptake in *S. cerevisiae* mutant strains deleted in essential metal transporters. Yeast strains, *ftr1*Δ (A), *mid1*Δ (B), *cch1*Δ (C), *ctr1*Δ (D), *fet3*Δ (E), *cot1*Δ (F), *smf1*Δ (G) and their wild type counterpart (BY4742 strain) were grown at late-log phase in YD medium and then suspended at a concentration of 2 × 10^8^ cells/ml in 10 mM MES, pH 5.5 buffer containing 20 mM glucose. After a 15 min preincubation at 30°C, 10 μM uranyl nitrate was added and cells were further incubated at the same temperature. At the time indicated, 1 ml aliquots of the cell suspensions were taken and processed as detailed in the Materials and Methods section. U present in cell pellets washed twice with 10 mM sodium carbonate was quantified by ICP-MS. Bars = SD; n = 2 to 6 independent experiments, as indicated. Statistical significance (mutant versus wild-type) was determined using a non-parametric Dunnett’s test with *P* < 0.001 (*).

Finally, the expression from a low copy plasmid, of wild-type copy of *MID1* gene under the control of its own promotor in the corresponding mutant strain restored its capacity to incorporate U to wild-type levels (Figure 7), confirming the role of the Cch1/Mid1 calcium channel in U transport in yeast. We were unable to perform reliable U uptake experiments using the *ftr1*Δ mutant bearing high copy number plasmids encoding Ftr1 and Fet3 proteins. Indeed, the co-expression of these recombinant proteins is controlled by iron deficiency (Stearman et al., 1996) and transformed cells have difficulties to grow properly on synthetic medium deprived of iron, as compared to control cells.

**Figure 7:**
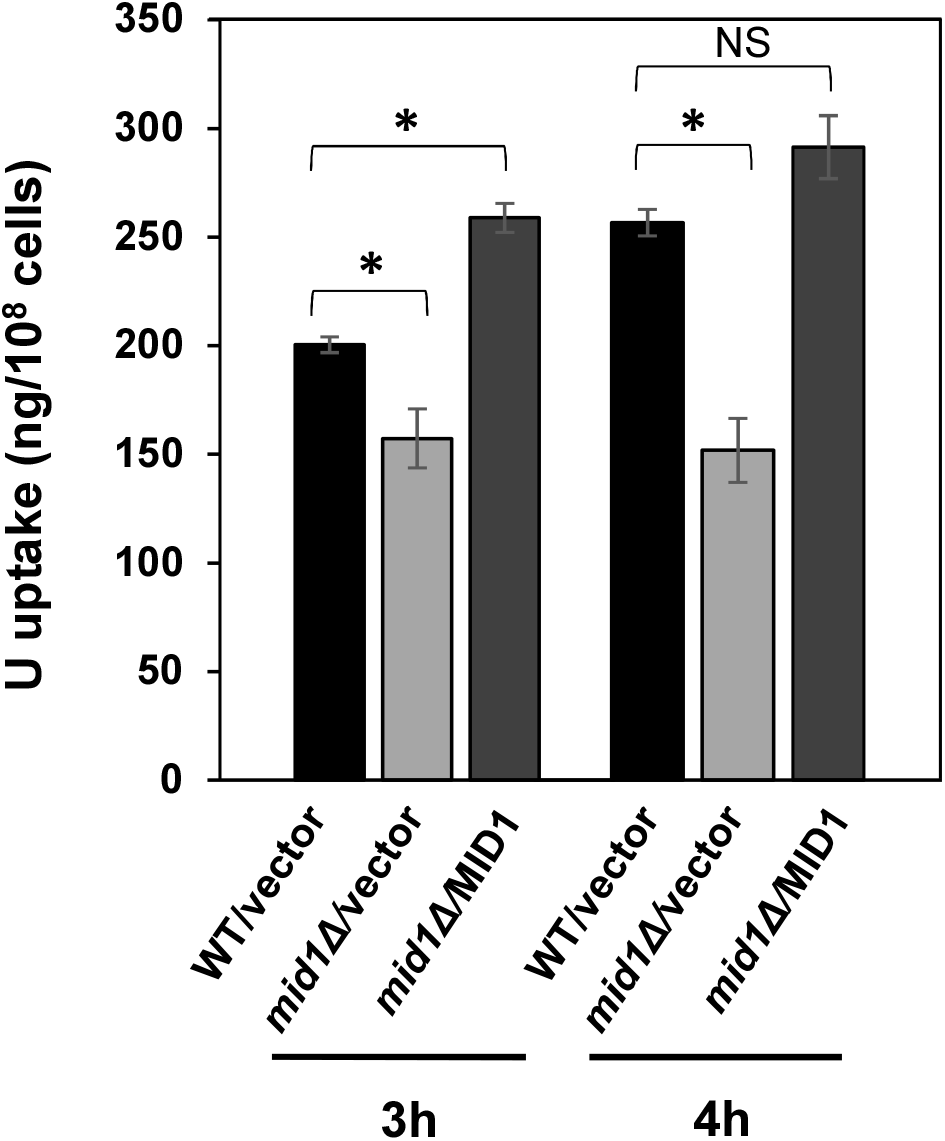
Contribution of the calcium channel Cch1/Mid1 to U entry in *S. cerevisiae* BY4742 strain. The yeast *mid1*Δ mutant as well as the wild-type parental BY4742 strain (WT) were transformed with either the empty low copy number plasmid pRS316 (vector; WT/vector and *mid1*Δ/vector) or YCpMID1 low copy number plasmid (MID1; *mid1*Δ/MID1) encoding the yeast *MID1* gene under the control of its own promoter. Freshly transformed cells were grown at late-log phase in SD/Ca100, plus amino acids and adenine, minus URA medium and then suspended at a concentration of 2 × 10^8^ cells/ml in 10 mM MES, pH 5.5 buffer containing 20 mM glucose. After a 15 min preincubation at 30°C, 10 μM uranyl nitrate was added and cells were further incubated at the same temperature. At the time indicated, 1 ml aliquots of the cell suspensions were taken and processed as detailed in the Materials and Methods section. U present in cell pellets washed twice with 10 mM sodium carbonate was quantified by ICP-MS. Error bars = SD; n = 3 independent transformants. Statistical significance was determined using a non-parametric Dunnett’s test with *P <* 0.001 (*). NS = not significant.

## Discussion

In this study, we identified, for the first time, calcium and iron assimilation pathways as important routes for U uptake into a living cell, using *S. cerevisiae* as a model. This was made possible, as discussed below, thanks to the development of an efficient procedure for measuring U transport, under conditions that favor U uptake and that allow to discriminate the uranyl fraction simply adsorbed on the cells from the intracellular fraction.

### Living yeast *S. cerevisiae* is able to accumulate U

Over the last four decades, the interactions of the uranyl ion with microorganisms, and particularly the yeast *S. cerevisiae*, have been extensively studied because of their potential use to remediate contaminated waste streams and groundwater (Kolhe et al., 2020; Lu et al., 2013; Nakajima and Tsuruta, 2002; Strandberg et al., 1981). Non-pathogenic and easy obtained at low cost from fermentation industry, *S. cerevisiae* was shown to display a high capacity of U biosorption, a biosorption which consists essentially of adsorption on the cell wall in the experimental conditions used (Chen et al., 2020; Ohnuki et al., 2005; Volesky and May-Phillips, 1995; Zhang et al., 2020). The most recent investigations proposed a three-step process for biomineralization of U on *S. cerevisiae* cell surface: surface adsorption, amorphous precipitation, and crystallization (Shen et al., 2018; Zheng et al., 2018). U biosorption conditions described so far, using complex media or unbuffered nutrients-free solutions, living or dead cell biomass exposed to elevated U concentrations generally higher than 0.4 mM were optimized for massive U precipitation and/or biomineralization on cell surface. In these conditions, U internalization was rarely evoked and, if so, it was mainly explained by the loss of the cell wall and plasma membrane integrity (Ohnuki et al., 2005; Strandberg et al., 1981; Volesky and May-Phillips, 1995).

In our work, U uptake experiments were performed on living cells exposed to low U concentrations (<100 μM) to preserve cell viability and to limit U precipitation on cell surface, in MES buffer adjusted to pH 5.5, which favors UO_2_^2+^, the bioavailable form of U. This buffer was chosen for its low to negligible metal-chelating propensity (Good et al., 1966) and to prevent pH neutralization of the uptake medium caused by gradual inorganic phosphate and ammonium ions release from inside the yeast cells upon U exposure (Shen et al., 2018; Zheng et al., 2018), minimizing uranyl phosphate precipitation on cell surface. In these conditions, U adsorption on cell surface was extremely rapid, around 80 % of U present in the medium being adsorbed in a few min. This observation compares well with data from literature showing equilibrium reached within 1 to 30 min according to experimental conditions (Faghihian and Peyvandi, 2012; Lu et al., 2013; Nakajima and Tsuruta, 2002; Popa et al., 2003; Volesky and May-Phillips, 1995). To distinguish U externally adsorbed on cell surface from U incorporated within the cell, we used two washings with carbonate, one of the highest affinity ligands for U (Kitano and Oomori, 1971; Poston et al., 1984). In our hands, two washing steps with 10 mM sodium carbonate allowed complete elimination of U adsorbed on cell surface, allowing to unmask U incorporated within the cell. This high washing efficiency was determinant for accurate measurement of uptake kinetics.

### Efficient U uptake into *S. cerevisiae* cells depends on the availability of a metabolizable substrate and necessitates a functional respiratory chain

In this work, we showed that an efficient U uptake requires an active metabolism. Indeed, net transport was almost null at 4°C regardless of the presence or absence of externally added glucose in the uptake medium, and low at 30°C in its absence, but it was highly stimulated in its presence. Interestingly, at low U concentrations (up to 3 μM U) glucose-independent U transport was significant, suggesting passive diffusion contribution to U uptake under these conditions. At 10 μM U, glucose-independent U uptake was almost negligible. At 30°C and in the presence of glucose, U uptake was a saturable process, initial rates of U uptake being saturable at concentrations above 50 μM. In *S. cerevisiae* glucose uptake is mainly performed by facilitated diffusion and is mediated by various uniporters and channels (Boles, 1997; Kruckeberg, 1996; Reifenberger et al., 1997; Wijsman et al., 2019). Since U and glucose are known to form complexes under certain conditions (Steudtner et al., 2010) the hypothesis of a co-transport through hexose transporters could not be excluded at this stage. To test this hypothesis, we used glycerol, which is not able to form complexes with U, as an alternative source of carbon in U uptake experiments. Similar stimulation of U uptake was observed under these conditions, ruling out the possibility of U transport through hexose transporters. The higher rates of U incorporation in the absence of any energy source in uptake medium, when yeasts were initially grown in medium containing glycerol instead of glucose (YG vs YD medium) (Figure 4B), might be explained by some existing internal storage pool of glycerol. In contrast, in yeast exposed to fermentable sugars such as glucose, despite high rates of uptake, little or no sugar can normally be detected within the cell, the glycolytic system removing the sugars as fast as they enter (Klein et al., 2017).

Finally, the use of sodium azide that blocks cell respiration by inhibiting the activity of mitochondrial cytochrome c oxidase-respiratory chain complex (Keilin, 1936; Leary et al., 2002), almost completely abolished U uptake in *S. cerevisiae* cells, whatever the metabolizable substrate source present (glucose or glycerol), demonstrating the importance of a functional respiratory chain in this process. The unaltered viability (cell permeability) of azide-treated cells compared to the control precluded the scenario that the very low U level in these cells resulted from cell death and the release of U to the medium (Supplemental Figure 1B).

### Iron, calcium and copper pathways are involved in the absorption of U in *S cerevisiae*

A combination of competition experiments with essential metals and analysis of mutant strains deleted in genes coding for metal importers allowed us to identify calcium, iron and copper pathways as potential routes for U uptake in yeast.

#### Uptake of U through the calcium assimilation pathway

In this study, we showed that in *S. cerevisiae* calcium and U share a common route for the entry into the cell. U(VI) under its UO_2_^2+^ cation form resembles Ca^2+^. Uranyl coordination properties having some similarities with those of calcium, various calcium-binding proteins from different organisms, including Human, were found to interact strongly with U, among which calmodulin (Brulfert et al., 2016; Le Clainche and Vita, 2006; Pardoux et al., 2012) and osteopontin (Creff et al., 2019; Qi et al., 2014; Safi et al., 2013). In biological systems, U and calcium biosorption relationships are also well documented. In *S cerevisiae*, Ca^2+^ was shown to affect both the initial rate of U biosorption and total amount of U bound to the cell wall (Strandberg et al., 1981). In plants, it was also found that U accumulation in plants was directly related to mobile calcium content in soil (Ratnikov et al., 2020). The interference of U with calcium uptake and homeostasis has been also reported in numerous studies (El Hayek et al., 2018; Moll et al., 2020; Rajabi et al., 2021; Straczek et al., 2009; Tawussi et al., 2017; Vanhoudt et al., 2010). The observed increase in cytoplasmic calcium reported in some cases could represent a cellular defense mechanism in response to the presence of U (White and Broadley, 2003). Within the cell, U(VI) under its UO_2_^2+^ cation form can replace Ca^2+^ and Mg^2+^, which can lead to structural changes to cell membranes, enzyme inactivation, and damage to RNA and DNA (Saenen et al., 2013). Despite all these observations, no direct evidence for the uptake of U by the calcium assimilation pathway was provided so far.

In *S. cerevisiae*, Mid1 (positive regulatory protein) and Cch1 (pore-forming subunit) are the two mandatory components of the high affinity voltage-gated calcium channel (Fischer et al., 1997; Hong et al., 2013; Iida et al., 1994; Iida et al., 2004; Iida et al., 2017; Iida et al., 2007). The Mid1/Cch1 channel becomes activated by various stimuli including depolarization, depletion of secretory calcium (Locke et al., 2000), pheromone stimulation (Paidhungat and Garrett, 1997), hypotonic shock (Batiza et al., 1996), and alkaline stress (Viladevall et al., 2004) or metallic stress (Gardarin et al., 2010). Interestingly, the Mid1/Cch1 calcium channel was previously found to contribute also to Cd^2+^ uptake in yeast (Gardarin et al., 2010).

In our study, we showed that calcium inhibits U uptake in *S. cerevisiae* in a concentration-dependent manner, suggesting a common entry pathway for these two cations. This hypothesis is supported by the fact that UO_2_^2+^ and Ca^2+^ are both divalent cations of similar atomic radius (around 180 pm) and ionic radius (around 80-100 pm). Indeed, the size and shape of the species handled by channels need to match the size of the ion selectivity filter cavity for a proper transport (Gouaux and Mackinnon, 2005). Experimentally, this hypothesis was also supported by the fact that the mutant strains *mid1*Δ and *cch1*Δ had similar phenotypes *i.e.*, low U uptake activity and accumulation, consistent with their contribution to a single functional complex, and that the expression of *MID1* gene into the *mid1*Δ strain restored wild-type levels of U uptake rates (Figure 6B,C). Thus, it is interesting to note the parallel between the kinetics of incorporation of U in calcium import mutants in our study and those for calcium in equivalent mutant strains (Iida et al., 1994; Nakagawa et al., 2007). Together, these observations demonstrated that the main calcium import pathway in yeast is involved in U transport.

The incomplete inhibition of U uptake in *mid1*Δ and *cch1*Δ mutant strains or in wild-type strain challenged with excess calcium suggests that other pathways are involved. A similar conclusion could be drawn for calcium entry in yeast cells (Cui et al., 2009; Iida et al., 1994).

#### Uptake of U through the iron assimilation pathways

Our study also suggests that U might enter the cell through iron transport systems. As for calcium, U and iron crosstalk is well documented in biological systems, particularly in plants where it was found that U interferes with iron homeostasis (Berthet et al., 2018; Doustaly et al., 2014; Sarthou et al., 2020; Vanhoudt et al., 2011). Also, several ferric iron-binding proteins were found to be able to bind U with high affinity, such as human transferrin, involved in iron transport to the bloodstream (Vidaud et al., 2007) and ferritin, an iron storage protein, from the archea *Purrococus furiosus*, the crayfish *Procambarus clarkia* or the zebrafish *Danio rerio* (Cvetkovic et al., 2010; Eb-Levadoux et al., 2017; Xu et al., 2014).

In the present study, we provided the first experimental evidence that the iron pathway can directly contribute to U uptake in yeast. Indeed, we showed that a mutant strain lacking the ferric iron permease Ftr1 was significantly affected in U uptake. The presence of functional copper-dependent ferrous oxidase Fet3 being required for the maturation of Ftr1 and iron transport across the plasma membrane (De Silva et al., 1995; Hassett et al., 1998a; Stearman et al., 1996) (Figure 8), the rather normal U uptake observed in the mutant strain *fet3*Δ was unexpected (Figure 6E). However, it was shown that yeast can compensate for the lack of Fet3 by overexpressing Fet4, a low affinity, low specificity Fe(II) ion transporter. This compensation was not observed in the mutant strain *ftr1*Δ (Li and Kaplan, 1998). Thus, in Fet3 defective mutants, U might enter the cell through the Fet4 pathways. In Ftr1 defective mutants, iron enter the cell mainly *via* Smf1, a broad-spectrum low affinity, low saturable divalent cation transporter capable of transporting Mn(II), Zn(II) and Fe(II) (Cohen et al., 2000; Portnoy and Culotta, 2003). Since Smf1 does not seem to be involved in U transport (Figure 6G), this would explain the deficit in U incorporation observed in the *ftr1*Δ mutant. These observations are consistent with the results of competition assays showing that a Fe(II) concentration about 10 times higher than that of Fe(III) was necessary to affect U uptake (Figure 5B,D). Taken together, our results support the idea that *S. cerevisiae* can incorporate U, through the Ftr1/Fet3 high affinity iron transport system.

**Figure 8:**
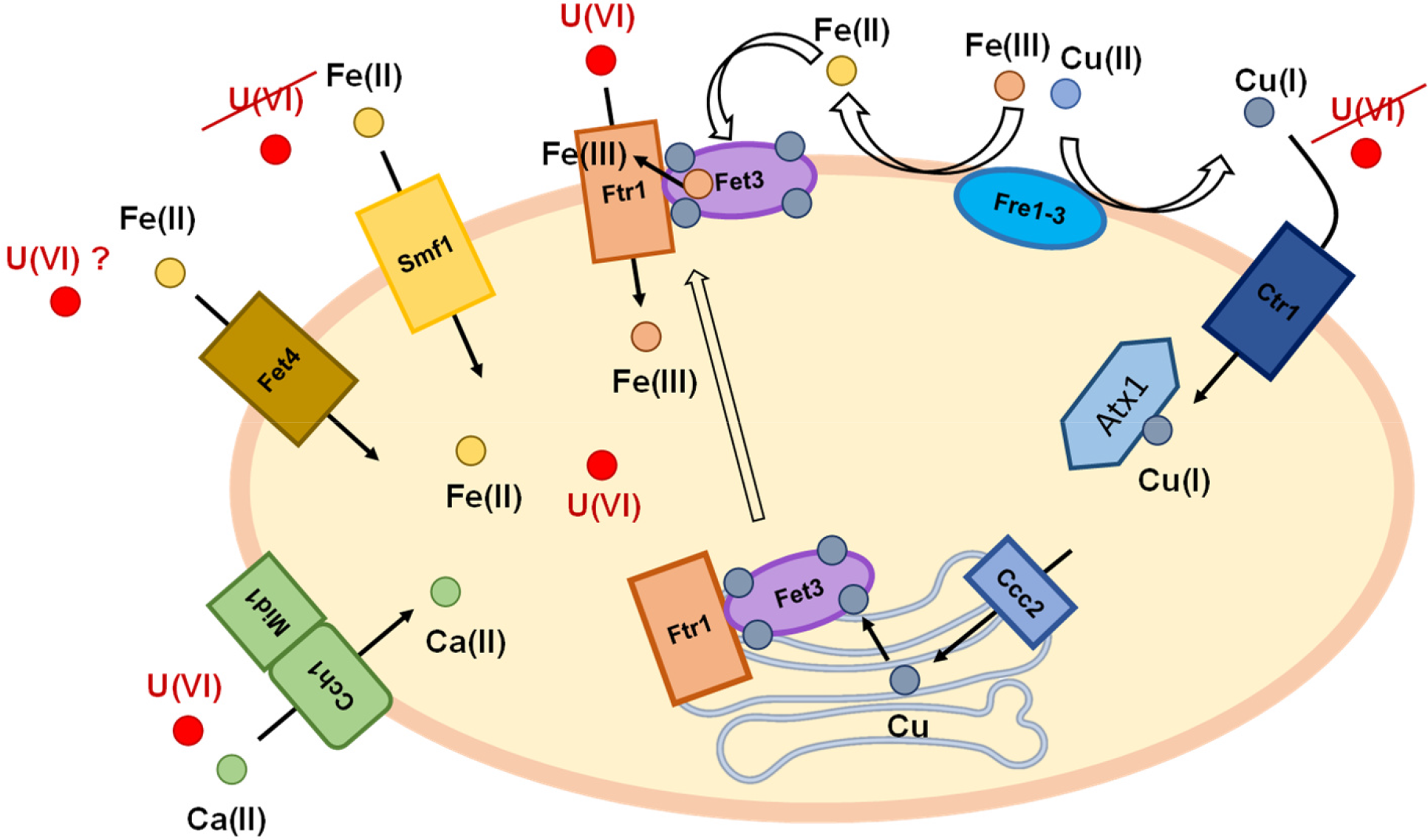
Recapitulative scheme of U entry routes in *S. cerevisiae*. Our data support the assertion that the high affinity Mid1/Cch1 calcium channel and the high affinity Ftr1/Fet3 iron transport complex promote U uptake in *S. cerevisiae*. Mid1/Cch1: the high affinity calcium channel; Fre1-3: Fre1, Fre2 and Fre3 reductases reduce the Cu(II) to Cu(I) and Fe(III) to Fe(II); Ctr1: the high affinity Cu(I) transporter; Atx1 chaperonin allows copper supply to the Golgi device in concert with Ccc2 transporter. In the path of secretion, copper is incorporated to various metalloproteins including multi-copper ferrous oxidase Fet3. Once copper incorporated in Fet3, the latter is secreted on the cell surface together with the permease Ftr1 with which Fet3 is physically associated, promoting the transport of iron inside the cell (Ftr1/Fet3, the high affinity iron transport complex). Fet4: low affinity and low specificity Fe(II) ion transporter; Smf1: broad-spectrum low affinity and low saturable divalent cation transporter.

#### Uptake of U under copper excess

The negative impact of copper on U uptake by yeast was unexpected. Indeed, the imported form of copper Cu(I) in yeast has very different chemical properties compared to U, notably in terms of protein ligands as deduced from Pearson’s (Pearson, 1963) classification. Accordingly, the mutant strain *ctr1*Δ lacking the high affinity Cu(I) transporter Ctr1 was not affected in U uptake (Figure 6D). We can exclude that Ctr3, another high affinity copper transporter, compensate for the lack of Ctr1 since the *Ctr3* gene is inactivated in most laboratory strains by a TY2 transposon, including S288C, the parental strain of BY4247 used in our study (Knight et al., 1996). The inhibitory effect of copper on U uptake might be indirect. For example, it was shown that copper stress induces broad and prolonged rise in cytosolic calcium through vacuolar and endoplasmic reticulum stored calcium release (Jo et al., 2008; Ruta et al., 2016) leading to downregulation of the Mid1/Cch1 calcium influx channel (Hong et al., 2010). Thus, in such a scenario, copper effect on U uptake would be explained by a crosstalk between copper and calcium homeostasis. Similarly, as described above, copper through its interaction with Fet3 is required for high affinity iron transport in yeast (Figure 8) (Dancis et al., 1994; Hassett et al., 1998b). It was shown that exposure to high copper concentrations could indirectly affect the iron transport machinery by compromising copper loading on Fet3 and subsequent translocation of the complex to the plasma membrane (Jo et al., 2008). In this scenario, exposure to copper would indirectly affect U uptake. Finally, at the concentration used in this study, copper slightly affected cell viability (15% of cells were dead after 3h incubation in the presence of 100 μM copper; Supplemental Figure 1C). Thus, we cannot exclude that inhibition of U uptake by copper was also related to the pleiotropic toxic effects of the latter on yeast metabolism (Shanmuganathan et al., 2004).

## Conclusion

In the present work, we have developed an accurate and reliable assay to elucidate U uptake routes in *S. cerevisiae*. Taken together, our data combining energy source requirement experiments, competition assays and mutant analyses, showed that U uptake probably operates through different transport systems, among which the calcium channel Mid1/Cch1 and the iron transport complex Ftr1/Fet3. This is to our knowledge, the first identification of U uptake routes in a living cell. Thus, identifying U-transporters in yeast, represents an important milestone in the fields of radionuclide toxicology and bioremediation. Particularly, our discoveries make possible rapid and systematic *in levuro* functional testing of candidate U transporters of many organisms, regardless of their phylogenetic origins. This can be particularly well suited to identify transporters from multicellular organisms, like higher plants or mammals, prior more complex and time-consuming characterization in their natural context. Validation of yeast-based hypotheses in these organisms would provide new insights into the chemical toxicity of U in higher eukaryotes and is a prerequisite for a future rational management of U in polluted soils, in the food chain, and in human health preservation.

## CRediT authorship contribution statement

**Revel B.**: Methodology, Investigation, Formal analysis. **Catty P.**: Conceptualization; Methodology, Investigation, Formal analysis, Resources, Writing – Review & Editing. **Ravanel S.**: Conceptualization, Formal analysis, Resources, Writing – Review & Editing; **Bourguignon J.**: Conceptualization, Supervision, Methodology, Formal analysis, Resources, Writing – Review & Editing, Funding acquisition. **Alban C.**: Conceptualization, Supervision, Methodology, Investigation, Formal analysis, Resources, Writing – Original draft, Writing – Review & Editing.

## Declaration of Competing Interest

The authors declare that they have no known competing financial interests or personal relationships that could have appeared to influence the work reported in this paper.

## Acknowledgements

This work was funded by grants from the Agence Nationale de la Recherche (ANR-17-CE34-0007, GreenU Project; ANR-17-EURE-0003, CBH-EUR-GS). The PhD fellowship to B.R. was funded by the CEA (CFR Grant). We gratefully acknowledge Pr Hidetoshi Iida and Dr Robert Stearman for providing us with plasmid constructions used in this study.

## Appendix A. Supplementary Material

## Supporting information

**Figure S1:**
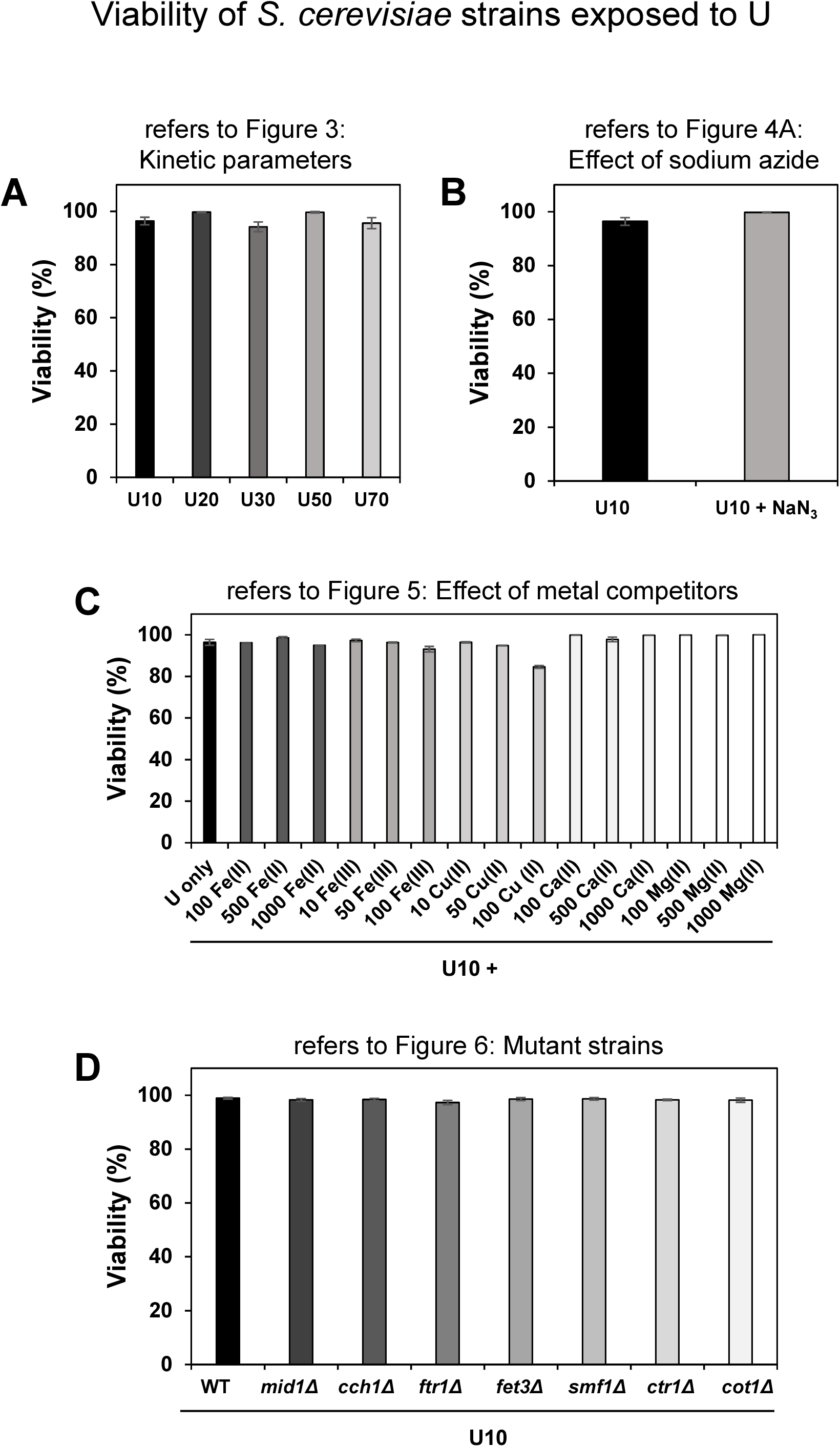
Viability of *S. cerevisiae* strains in the course of the experiments performed in this study. Viability of yeast cell suspensions (Parental BY4742 wild-type strain and the related isogenic mutant strains) was determined by the fluorescein diacetate/propidium iodide staining method using Yeast viability kit and the automated fluorescence cell counter LUNA-FL (Logos Biosystems) after 4h incubation of cells at 30°C in the presence of 20 mM glucose, uranyl nitrate at the indicated concentrations (in μM), and other molecules (as indicated) in 10 mM MES pH 5.5 buffer. The details of growth and assay conditions are given in the Materials and Methods section and in the legend of the figures of the corresponding experiments. (A) Viability of cells from experiments in Figure 3 (Kinetics parameters determination); (B) Viability of cells from experiments in Figure 4A (Effect of sodium azide); (C) Viability of cells from experiments in Figure 5 (Effect of metal competitors). Concentrations of competitors are given in μM from 10 to 1000; (D) Viability of cells from experiments in Figure 6 (Mutant strains deleted in essential metal transporters). The results from viable cell counts are shown as relative values (%) as compared to total counted cells. Error bars = SD; n = 2 to 6 independent experiments, as indicated in the legend of the referred figures.

## Notes

### Competing Interest Statement

The authors have declared no competing interest.

